# Rheumatoid Arthritis-associated IgG N-glycan agalactosylation diminishes neutrophilic inflammation by reducing FcγR binding and downstream signaling

**DOI:** 10.64898/2026.07.02.735866

**Authors:** Caroline Pumpe, Alfie Sanderson, Ballagh Forsyth, Jelena Šimunović, Yoshiki Narimatsu, Henrik Clausen, Gordan Lauc, Mark S. Cragg, Pierre Bruhns, Mohini Gray, Cécile Bénézech, Caroline Hayward, Sonja Vermeren

## Abstract

The IgG Fc chain carries a single N-linked glycan which may undergo changes. Increased agalactosylated N-glycans are associated with rheumatoid arthritis (RA) and regarded as pro-inflammatory. Dysregulated neutrophils can make important contributions to host tissue damage. In RA, immune complexes (ICs) that have precipitated onto synovial joint surfaces activate neutrophils via Fc receptors, promoting localised inflammation. We engineered recombinant human monoclonal IgG with agalactosylated or galactosylated N-glycans, generated immobilised ICs and stimulated healthy donor and RA patient blood-derived neutrophils, comparing reactive oxygen species (ROS) production as read-out of neutrophilic inflammation. Both healthy donor and RA patient neutrophils generated *less* ROS when stimulated with ICs made from agalactosylated IgG. Mechanistically this was due to poorer binding of agalactosylated ICs to neutrophil FcγRs, causing lower activation of Akt and p38 MAPK. Both are required for immobilised IC-mediated stimulation of the neutrophil NADPH oxidase. Taken together, this suggests that disease-associated, agalactosylated IgG does not in fact promote inflammation and host tissue injury, at least not by acting on neutrophils. We propose that rather than promoting inflammation, agalactosylated IgG N-glycans that accompany inflammatory disease may arise as part of a compensatory mechanism that is aimed at reducing excessive inflammation and host tissue injury.

**Graphical Abstract:** 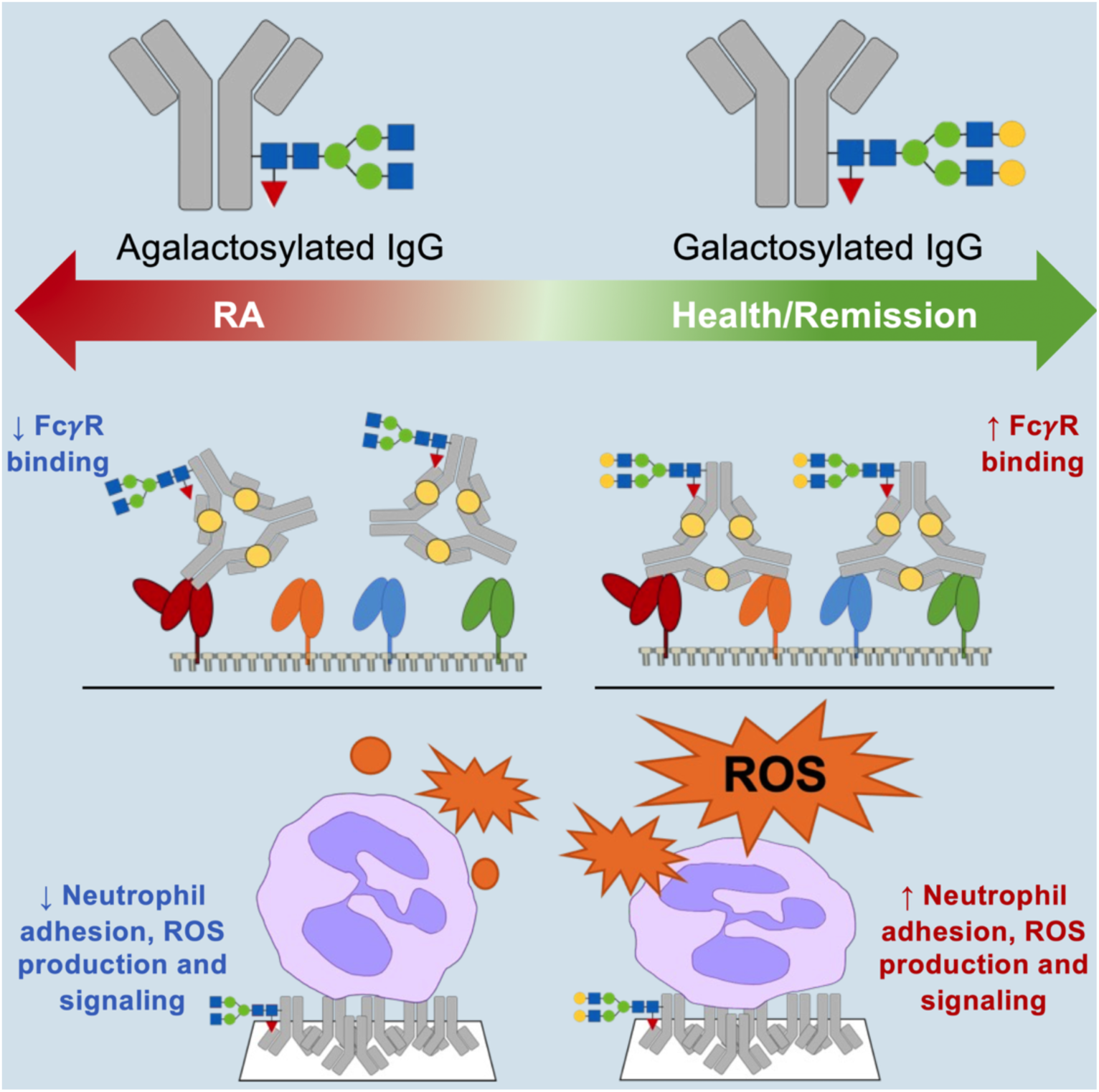

## Introduction

IgG, the most abundant antibody in human serum, is characterised by a single conserved Fc region-localised N-linked glycosylation site, asparagine 297. IgG glycosylation is mediated by glycosyltransferases, which are thought to be controlled epigenetically. The IgG N297-glycan consist of a bifurcated core, made up of two acetylglucosamine and three mannose residues. The core is typically fucosylated, and it may carry galactose and sialic acid residues. In disease, and old age, IgG glycosylation may undergo dynamic changes. Rheumatoid arthritis (RA) is an autoimmune disease primarily affecting synovial joints, in which autoantibodies form immune complexes (ICs) that promote inflammation (1, 2). In RA, both autoantibody and total serum IgG galactosylation (but not fucosylation) are lowered (3, 4). This altered glycosylation pattern has also been observed in other autoimmune diseases, including, systemic lupus erythematosus, multiple sclerosis, Crohn’s disease, primary biliary cholangitis and ulcerative colitis (5). In contrast, in youth, pregnancy and in RA patients in remission, the balance is shifted towards galactosylated IgG N-glycans. IgG carrying agalactosylated N-glycans is widely regarded as pro-inflammatory or even pathogenic, whereas galactosylated (and sialylated) IgG are assumed to be anti-inflammatory (6–9). Yet, whether and how differential antibody galactosylation and sialylation affects neutrophils in RA and elsewhere remains poorly understood.

Neutrophils, the most abundant circulating leukocytes in humans, hold a crucial function in host defence. If dysregulated, they can make important contributions to host tissue injury (10). ICs made up of (auto)antibody and (auto)antigen may precipitate on biological surfaces, e.g. in the synovial joint, contribute to the recruitment of neutrophils, and activate neutrophils by binding to their FcγRs (11). This results in neutrophil adhesion, frustrated phagocytosis and the release of reactive oxygen species (ROS) and cytotoxic degranulation products directly onto the biological surface on which the ICs are immobilised, promoting host tissue damage (12).

We set out to identify whether, and how, agalactosylated IgG promotes neutrophilic inflammation. We used genetically engineered Chinese hamster ovary (CHO) cells to generate anti-HSA designer IgG with defined glycans (13). We plated human neutrophils onto ICs made with agalactosylated or galactosylated IgG and observed *less* adhesion and ROS by neutrophils that had been plated onto agalactosylated ICs. Binding studies established that ICs made up of agalactosylated IgG bound less well to CD64 (FcγRI), CD32A (FcγRIIA), CD32B (FcγRIIB) and CD16B (FcγRIIIB). Immobilised ICs made up of agalactosylated IgG were less able to activate neutrophil signaling downstream of FcγR engagement, which is required for full activation of the phagocyte oxidase. We conclude that agalactosylated IgG is not inflammatory per se; rather it may be generated in order to limit excessive inflammation and host tissue damage.

## Results

We set out to test whether and how IgG carrying agalactosylated N-glycans, that are inflammation-associated, trigger more neutrophilic inflammation than health-associated IgG carrying galactosylated glycans. We took a reductionist approach, analysing how freshly isolated, unprimed peripheral blood-derived neutrophils reacted to being stimulated with immobilised ICs generated from human serum albumin (HSA) and recombinant, monoclonal anti-HSA IgG antibodies. To do this we generated recombinant, human anti-HSA IgG1 species carrying inflammation-associated, agalactosylated (G0F) and health-associated, galactosylated glycans (G2F; Fig 1A) by expressing the same IgG1-encoding construct in two different, genetically engineered CHO cell lines (13, 14). Recombinant IgG was purified and N-glycans were analyzed by lectin overlay (Fig 1B) and also by ultra-high performance liquid chromatography-mass spectrometry (Fig 1C). This confirmed that the majority of agalactosylated IgG was agalactosylated, while the majority of galactosylated IgG was galactosylated.

**Figure 1.**
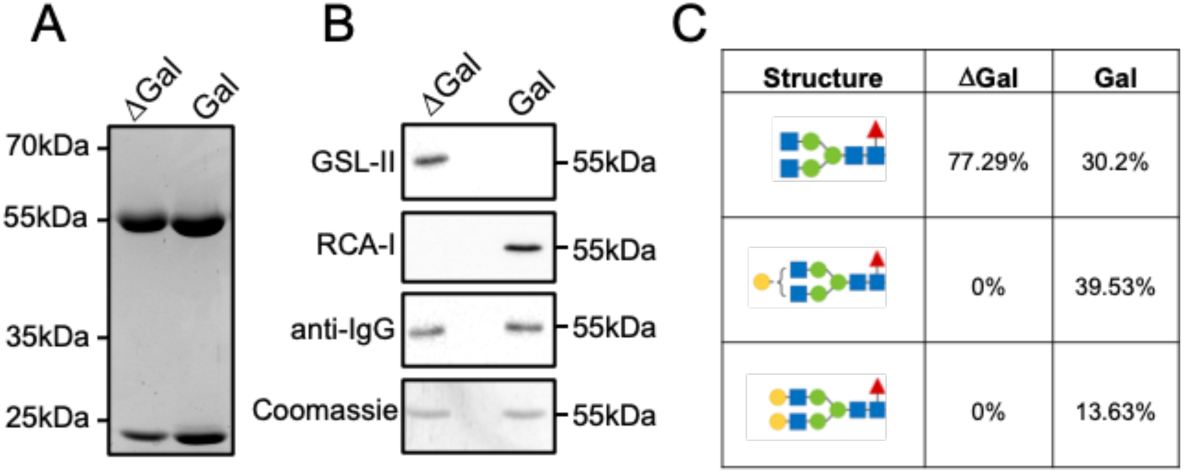
IgG1 with defined glycans produced in glycoengineered CHO cell lines. (A) A coomassie-stained protein gel of purified, recombinant agalactosylated (βGal) and galactosylated (Gal) anti-HSA IgG. (B) Validation of N-glycans by lectin overlay with Griffonia Simplicifolia lectin II (GSL-II), which recognizes the nonreducing terminal of α- or β-linked N-acetylglucosamine (GlcNAc) residues and with Ricinus communis agglutinin I lectin (RCA-I) which recognizes terminal β-galactose (Galβ) and Galβ-1-4GlcNAc residues. (C) Elucidation of the composition of released N-glycans by ultra-high performance liquid chromatography-mass spectrometry. Blue square, GlcNAc; red triangle, fucose; green circle, mannose; yellow circle, galactose. See Fig S1 for a full summary of all glycan traits.

### IgG1 N-glycans are sufficient to regulate FcγR-mediated neutrophil functions

We generated immobilised ICs made up of HSA and agalactosylated or galactosylated anti-HSA, respectively, and plated freshly isolated, unprimed human neutrophils onto the immobilised ICs. For this, neutrophils were obtained from healthy volunteers and RA patients recruited at an early arthritis clinic.

Contrary to expectation, we observed significantly *less* neutrophils from healthy donors adhered to immobilised ICs made with agalactosylated than galactosylated IgG. This pattern was also observed with neutrophils isolated from RA patient blood (Fig 2A,B). In contrast, neutrophil spreading and shape were not dramatically affected by the IgG glycan (Fig 2C,D).

**Fig 2.**
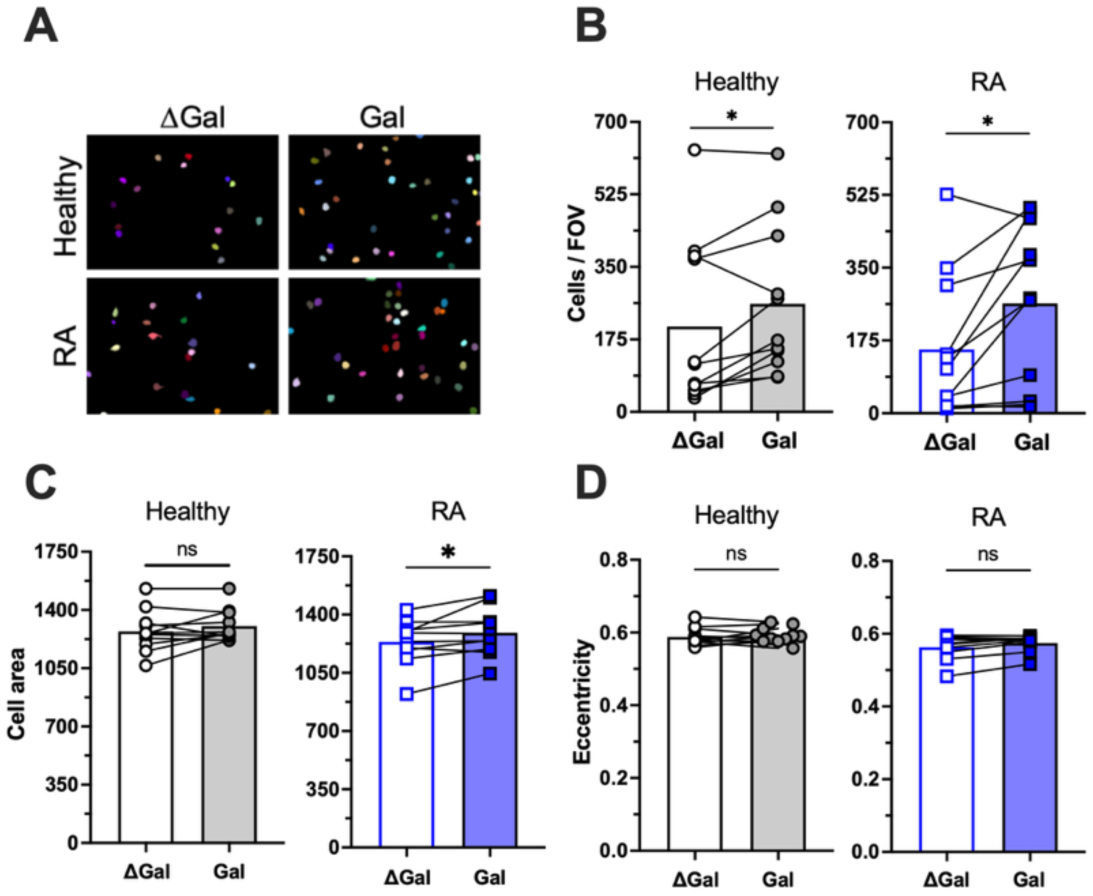
Neutrophils adhere better to ICs made from galactosylated, than agalactosylated IgG. Neutrophils were prepared from healthy donors and early RA patients and plated onto immobilised ICs made with galactosylated (Gal) or agalactosylated (ΔGal) IgG. Images of fixed cells were analyzed computationally. (A) Representative images. (B) Adhered cells per field of view, (C) neutrophil spreading (cell area) and (D) eccentricity are plotted. (B-D) Each symbol represents the average value obtained with neutrophils from one healthy donor or RA patient. Raw data were analyzed by Wilcoxon signed-rank test*. * p<0.05; ns not significant*.

As an indirect read-out of neutrophilic inflammation we also analyzed ROS production by neutrophils that had been plated onto immobilised ICs. In agreement with the adhesion experiments, we observed significantly *less* ROS production with healthy donor and RA patient neutrophils that had been plated onto agalactosylated versus galactosylated ICs (Fig 3A). Concurrently, however, we noted more ROS production by RA patient than healthy donor blood-derived neutrophils for each condition tested (Fig 3B). Hence, similar amounts of ROS were generated by RA neutrophils plated onto agalactosylated ICs (35609 ± 5229 mean ± SEM) as by healthy donor neutrophils plated onto galactosylated ICs (34876 ± 3785). In summary, the IgG Fc glycan was sufficient to direct the amplitude of neutrophil responses upon stimulation with ICs and this response pattern was conserved in healthy donor and RA patient neutrophils. These *in vitro* data suggest that agalactosylated IgG, although inflammation-associated, is not *per se* pro-inflammatory.

**Fig 3.**
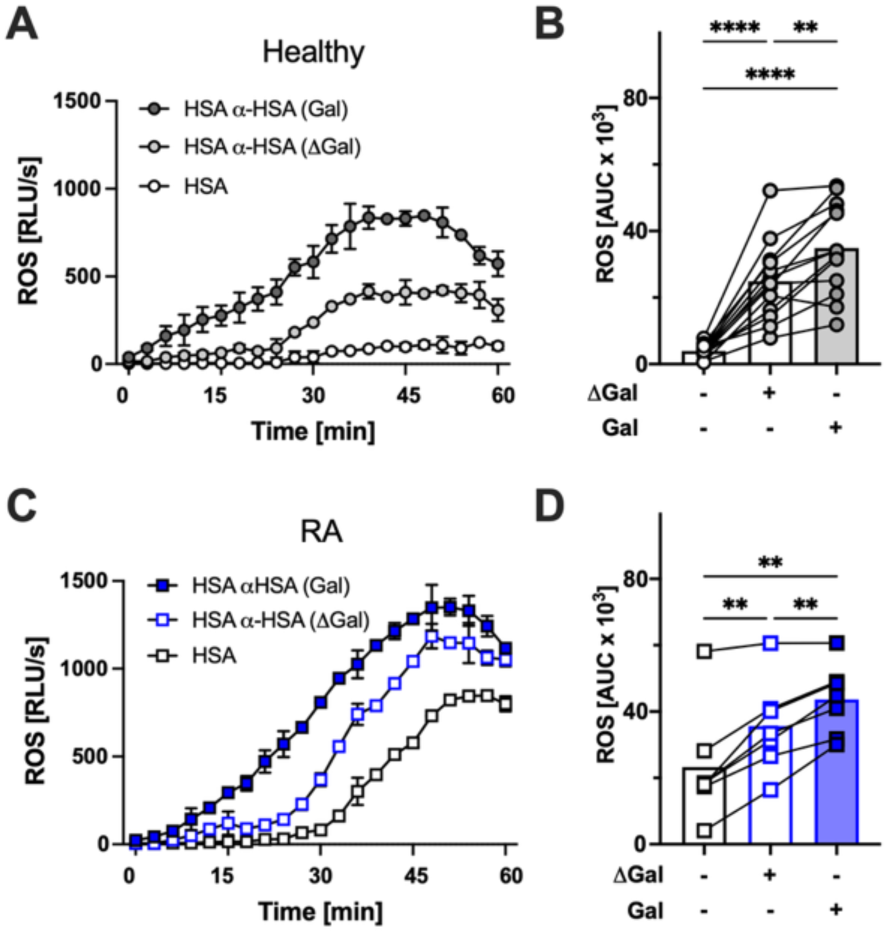
Neutrophils generate more ROS when plated onto galactosylated than agalactosylated ICs. Neutrophils that had been freshly prepared from (A,B) healthy donors and (C,D) early RA patients were plated onto immobilised ICs made with galactosylated (Gal) or agalactosylated (ΔGal) IgG. ROS production was measured in real-time. (A,C) Representative examples (mean ± range) and (B,D) integrated total ROS produced are plotted. (B,D) Each symbol represents one blood donor. Raw data were analyzed by one-way ANOVA with Tukey’s multiple comparison post-hoc test. ** p < 0.01; **** p < 0.0001.

### IgG N-glycans regulate FcγR binding

Human neutrophils express a range of FcγRs that bind IgG Fc. In addition to the highly abundant low affinity receptor CD16B, which is GPI-linked and not thought to transduce signals directly (15), they also constitutively express the abundant CD32A. They inducibly express the high affinity CD64 (16), and some express very low levels of the inhibitory CD32B (17). We performed flow cytometry to phenotype FcγR expression of freshly isolated neutrophils from healthy donors and RA patients (Fig 4), observing an increased percentage of CD64-positive RA patient neutrophils (Fig 4A). CD64 expression in neutrophils is induced by interferon-γ, and its increase in RA patient blood neutrophils is in agreement with a previous report (18). We did not observe differential expression of any FcγRs or indeed CD11b (Mac-1; Fig 4B-G) by neutrophils from healthy volunteers or RA patients.

**Fig 4.**
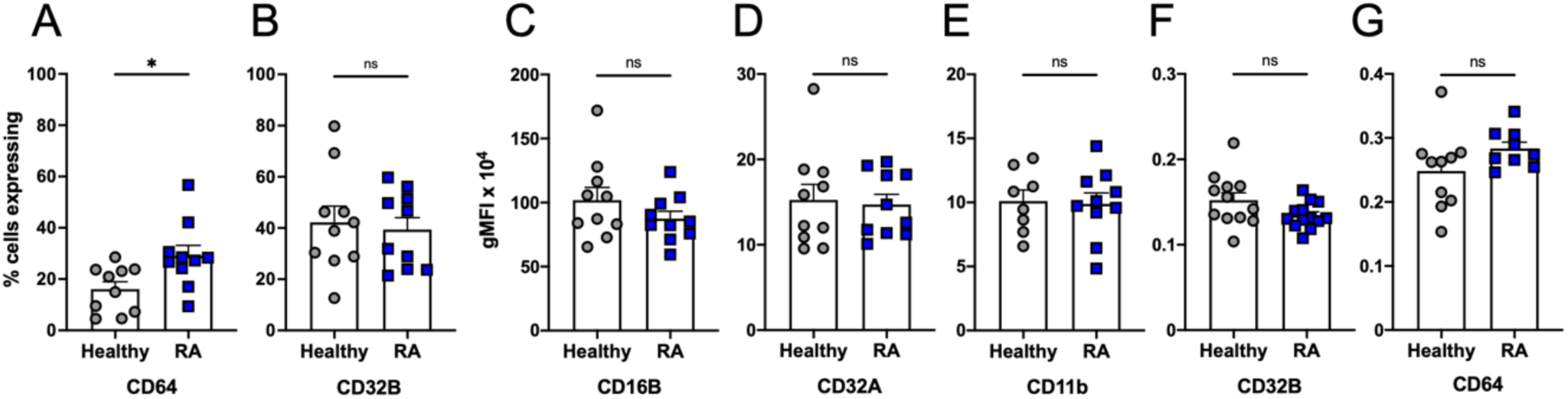
FcγR expression in neutrophils from healthy donors and RA patients. Neutrophils were isolated from peripheral blood, and receptor surface expression analyzed by flow cytometry as indicated. Each symbol represents one blood donor. Data were analyzed by Welch t-test; error bars show SEM. * p<0.05; ns, not significant.

Different single nucleotide polymorphisms (SNPs) are present in human CD32A [V158/F158] and CD16B [NA1/NA2] (19). We determined the SNPs carried by neutrophils used for this study but did not observe differences in SNP distribution between healthy donors and RA patients (not shown), which were predominantly of European ethnicity.

Since neutrophil FcγRs (with the exception of the lowly expressed FcγRI) are low affinity receptors, we analyzed binding of PE-labelled, small, defined ICs to heterologously expressed FcγRs on CHO cells by flow cytometry (20) (Fig S2) to test whether IgG (a)galactosylation affects receptor binding. We observed significantly *weaker* binding of agalactosylated compared to galactosylated IgG1 to CD16B (NA1 and NA2 alleles), CD32A (R131 and H131 alleles), CD32B and also CD64 (Fig 5). This identified that in the context of ICs, the presence of galactose on the IgG1 Fc N-glycans was sufficient to determine differential FcγR binding.

**Fig 5.**
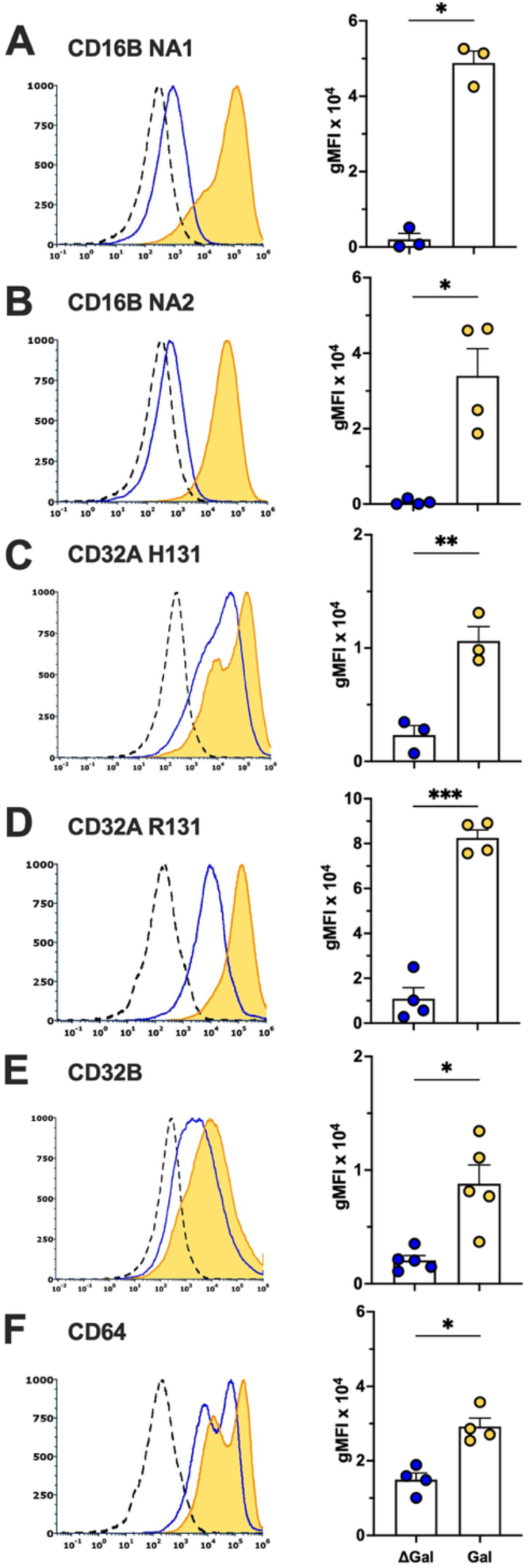
IgG N-glycans determine FcγR binding of ICs. Interaction of F(ab’)_2_-aggregated agalactosylated (ΔGal) and galactosylated (Gal) IgG with human FcγRs expressed on CHO cells was assessed by flow cytometry (A-F). Left, representative examples. Broken peaks, untransfected CHO binding to galactosylated ICs; continuous peaks, CHO cells expressing FcγR as indicated binding to agalactosylated (blue) and galactosylated (yellow) ICs. Right, summarised data from 3-5 independently conducted experiments. Each symbol represents the corrected gMFI obtained in one experiment. Blue, agalactosylated IgG; yellow, galactosylated IgG. Raw data were analyzed by two-tailed paired T test. Error bars show SEM * p<0.05; ** p<0.01; *** p<0.001.

### IgG1 N-glycans regulate signaling downstream of FcγR

FcγR signaling initiates multiple protein phosphorylation events in human neutrophils (12, 21, 22). We plated freshly isolated healthy donor human neutrophils onto immobilised ICs and analyzed the phosphorylation of Akt (an indirect read-out of PI3K activation), Erk and p38 MAPK using phosphospecific antibodies by western blot. We observed increased activation of Akt and p38 MAPK with neutrophils stimulated with galactosylated as opposed to agalactosylated ICs (Fig 6).

**Fig 6.**
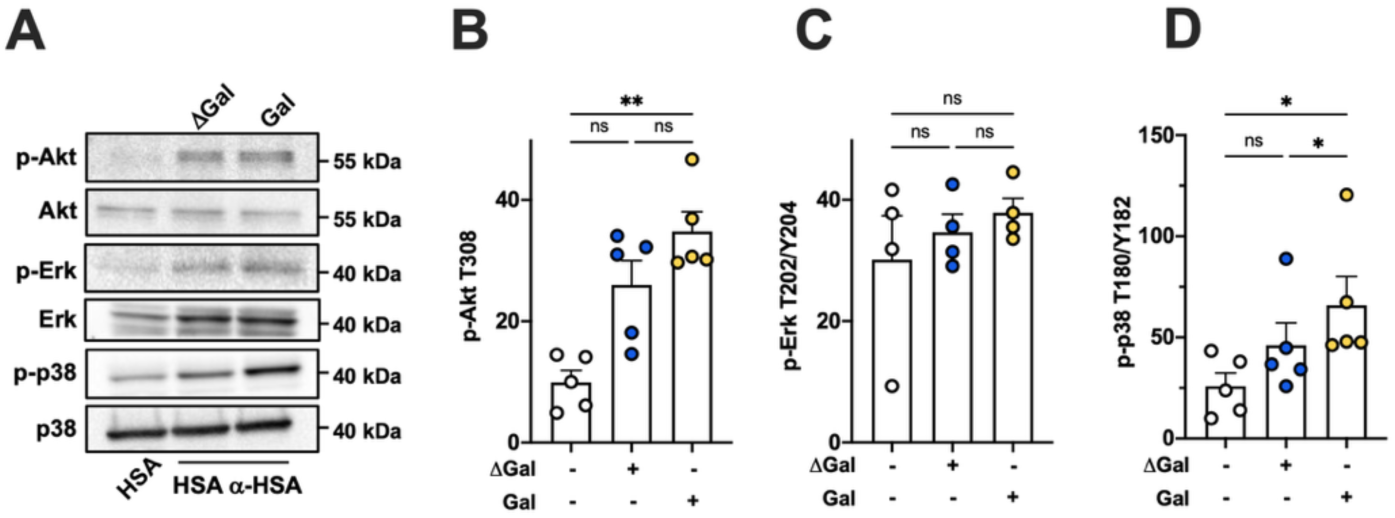
IgG glycan regulates neutrophil signaling. Neutrophils were stimulated by being plated onto immobilised ICs made with galactosylated (Gal) or agalactosylated (ΔGal) ICs and phosphorylation of Akt, Erk and p38 analyzed by western blot. (A) A representative example is presented alongside (B-D) integrated results from 4-5 separately conducted experiments. (B) Phospho-Akt, (C) phospho-Erk and (D) phospho-p38. (B-D) Each symbol represents one healthy donor or patient. Raw data were analyzed by one-way ANOVA with Tukey’s multiple comparison post-hoc test. Error bars show SEM. * p<0.05; ** p<0.01; ns, not significant.

### p38 MAPK regulates IC-induced ROS production

The phagocyte NADPH oxidase consists of multiple subunits, membrane located gp91^phox^ and p22^phox^, and the cytoplasmic p67^phox^, p47^phox^ and p40^phox^ subunits. To become active, the oxidase needs to assemble at a suitable membrane. This depends upon Rac-mediated translocation of p67^phox^ as well as phosphorylation of p47^phox^ relieving its autoinhibited conformation (23). PI3K regulates the NADPH oxidase via several mechanisms (24), including via activation of Rac2 by PtdIns(3,4,5)P_3_-dependent guanine nucleotide exchange factors. Erk and p38 MAPK were shown to be involved in NADPH oxidase in a context dependent fashion (25, 26), but their roles in immobilised IC-induced ROS remain unexplored. ROS production assays with human neutrophils plated onto immobilised ICs in the presence or absence of inhibitors targeting PI3K, Erk, p38 and the NADPH oxidase, identified regulatory functions for PI3K and p38 but not Erk activity (Fig 7). This suggests that by less efficiently promoting signaling downstream of FcγR engagement in neutrophils plated onto ICs, agalactosylated IgG leads to suboptimal activation of the NADPH oxidase.

**Fig 7.**
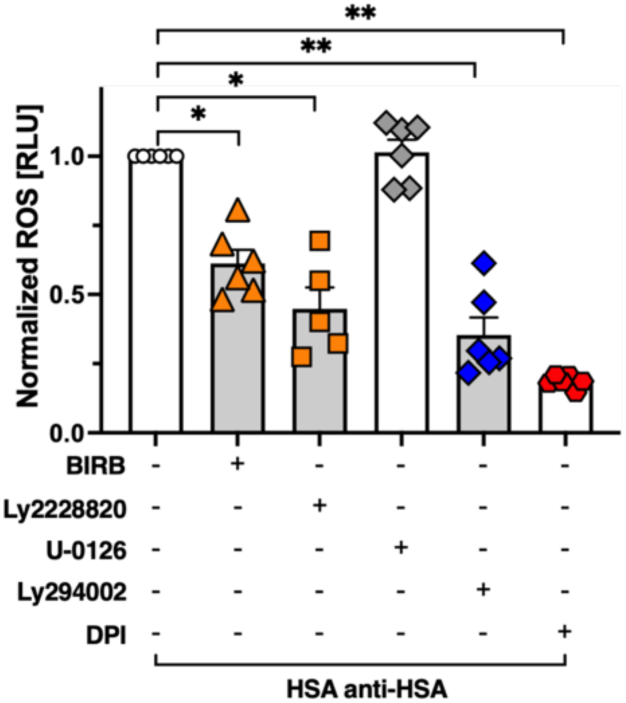
PI3K and p38 promote IC-stimulated ROS. Neutrophils that had or had not been treated with inhibitors targeting p38 (BIRB-796; LY2228820), Erk (U-0126), PI3K (LY294002) or the NADPH oxidase (DPI) were stimulated with ICs. ROS production was measured. Normalized results are presented; each symbol represents one healthy volunteer. Raw data were analyzed by one-way ANOVA with Tukey’s post-hoc test; comparisons relate to the vehicle-treated control; error bars show SEM. * p<0.05; ** p<0.01.

## Discussion

It was first described decades ago that antibody N-glycans differ in health and disease. In RA, agalactosylated IgG was repeatedly found to be elevated with agalactosylation correlating with disease activity and acquired prior to disease onset (27, 28). In contrast, periods of remission from RA were characterised by increased levels of galactosylated, and also of sialylated IgG, which are also elevated in the young, and in pregnancy (29). Elevated agalactosylated IgG was subsequently also observed in a range of other autoimmune diseases, as well as in old age, which is characterised by chronic low-grade inflammation (30). Perhaps largely because it is associated in this way with (chronic) inflammation, agalactosylated IgG is widely regarded as pro-inflammatory. In the lights of these correlations, our observations are unexpected.

Neutrophils are FcγR-expressing, highly abundant circulating leukocytes that are dysregulated in disease. ICs may precipitate on biological surfaces, where they act as powerful stimuli that promote neutrophil recruitment and elicit a range of neutrophil effector functions that include ROS production, and that together promote host tissue damage (2, 11). This places neutrophils as immune cells that are prone to being deeply affected by any effects of differential IgG glycosylation. To analyze the effect of IgG N-glycans on human neutrophils, we used ROS production, induced by immobilised ICs as an indirect read-out of neutrophilic inflammation. The monoclonal antibodies employed to generate the ICs differed only in their N-glycan composition, carrying either the health-associated galactosylated, or the disease-associated, agalactosylated glycoform. Rather than confirming that agalactosylation renders IgG inflammatory by better activating neutrophils, our observations suggest that galactosylated ICs are able to better activate neutrophils, eliciting more adhesion and increased ROS production. Mechanistically, this is because they better engage with and cross-link neutrophil FcγRs. These observations are likely to be pertinent to other FcγR-expressing effector cells that will also be affected by reduced receptor binding of agalactosylated IgG ICs.

We observed differential binding to heterologously expressed FcγRs on CHO cells with ICs generated from the different glycoforms, suggesting that our observations are due to a structural feature of the IgG N-glycan that is involved in mediating FcγR binding. This view is supported by NMR studies and computational modelling that both suggest IgG galactosylation stabilises the Fc region, promoting interactions with FcγRs (31). Other experimental evidence also supports our findings that glycosylation is critical for the interactions between IgG and its receptors. One previous study compared binding of ICs consisting of monoclonal full IgG, PNGaseF-treated IgG (where N-glycans are cleaved off entirely) or EndoS-treated IgG (where only the fucosylated N-acetylglucosamine and branching fucose remained), observing residual FcγR binding with the small glycan-containing, but not the aglycosylated ICs (32). It is moreover well established that IgG fucosylation regulates CD16A binding (20, 31, 33, 34), a function that is exploited in infectious diseases, where it renders CD16A-dependent cytotoxicity mediated by natural killer cells more powerful.

Unsurprisingly, improved FcγR engagement by galactosylated ICs resulted in increased activation of signaling intermediates including p38 MAPK and PI3K, which we showed to be required for optimal activation of the phagocyte NADPH oxidase in an IC-induced context. p38 MAPK had already been shown to be required for NADPH oxidase priming and activation in a context-dependent fashion (25, 26, 35).

We observed the same pattern, that galactosylated ICs induced more ROS, with neutrophils isolated from healthy donor and RA patient blood. Circulating neutrophils from the latter express more CD64, an IFN-γ inducible gene that was already known to be upregulated in RA neutrophils (16, 18). We noticed RA neutrophils produced more ROS in response to being plated onto immobilized ICs, an observation that is in keeping with previous observations with soluble stimuli (36). Interestingly, we noticed heightened ROS production by RA neutrophils even when they were plated onto HSA rather than immobilised ICs, suggestive of these cells being primed. Interestingly, RA patient neutrophils plated onto ICs made from agalactosylated IgG still produced as much ROS as those from healthy donors plated onto galactosylated IgG (Fig 3). This makes it tempting to speculate that IgG agalactosylation may arise as a compensatory mechanism aimed at curbing overexuberant inflammation and limiting host tissue damage.

In summary, the present work has uncovered a molecular mechanism by which antibody glycosylation directs neutrophilic inflammation, at least as read out by neutrophil adhesion and ROS generation. In doing so, our observations have identified that antibody agalactosylation is not in fact pro-inflammatory, at least not via neutrophils. Rather, our results suggest that ICs made from galactosylated antibodies are better able to elicit neutrophilic inflammation since they bind FcγRs more strongly. Interestingly, unrelated, recent analyses of complement binding and activation by different IgG1 glycoforms showed that complement activation is also significantly increased with galactosylated IgG glycoforms (31, 37, 38) (39). Given just how powerfully complement can kill, this observation is entirely in keeping with our conclusion that agalactosylated IgG is in fact hypoinflammatory.

## Materials and Methods

### Sex as a biological variable

Sex was not specifically considered as a biologically variable for the purpose of this study.

### Reagents

Reagents of the lowest possible endotoxin level were purchased from Merck Life Sciences (Glasgow, UK) unless indicated otherwise. Tissue culture reagents were obtained from Thermo Fisher Scientific (Loughborough, UK). Percoll was from GE Healthcare (Amersham, UK). Antibodies and inhibitors used in this study are summarised in tables S1-2.

### Human neutrophils

Peripheral venous blood was collected from male and female healthy volunteers and patients with written informed consent obtained before sample collection. Patients were recruited at an early arthritis clinic, where they were diagnosed and followed up for 12 months (Tables 1-2 for donor characteristics). Neutrophils were prepared as previously described (40). Neutrophil purity according to Diff-Quik (Thermo Fisher Scientific) stained cytospins exceeded 95%.

**Table 1.** Characteristics of healthy blood donors contributing to this study. . Age was captured in 10-year brackets.

| ***Total n (%)*** | ***Male n (%)*** | ***Female n (%)*** | ***Age Range*** | ***Median Age*** |
| --- | --- | --- | --- | --- |
| 22 (100%) | 8 (36%) | 14 (64%) | 21-60 yrs | 35-45 yrs |

**Table 2.** Characteristics of RA patients who donated blood used in this study. All RA patients were seropositive. 11 patients (85%) were newly diagnosed and 10 patients (77%) were treatment-naïve at the time of being included into the study.

| **Total**<br>**n (%)** | **Male**<br>**n (%)** | **Female**<br>**n (%)** | **Age**<br>**Range (yr)** | **Age (yr)**<br>**Median [IQR]** | **Non-smoker**<br>**n (%)** | **Smoker / Ex-**<br>**smoker n (%)** |
| --- | --- | --- | --- | --- | --- | --- |
| 13 (100%) | 4 (31%) | 9 (69%) | 33-78 yrs | 62 yrs<br>[59-66 yrs] | 3 (23%) | 10 (77%) |

### Cell Culture and production of recombinant human IgG

Cell lines were tested for microplasma contamination prior to being used for experiments. CHO-K1 cells expressing individual FcγR alleles (20, 41) were cultured in RPMI-1640 supplemented with Glutamine, 10% FBS (Cytiva, Little Chalfont, UK) as well as selection conditions as applicable for each cell line in a humidified atmosphere with 5% CO_2_ at 37°C. Previously described glycoengineered CHOZN GS-/- cells (14) (Merck Life Sciences) were cultured in serum-free EX-CELL CD CHO Fusion medium (Merck Life Sciences) supplemented with 4mM GlutaMAX (Thermo Fisher Scientific) and stably transfected with pVITRO1-ΔV-IgG1/κ (42) into which the clone 139 variable heavy and light chain specific for human HSA (43) had been inserted. Recombinant anti-HSA IgG was isolated using affinity chromatography using HiTrap protein L columns (Thermo Fisher Scientific) as per manufacturer’s instructions.

### Analysis of glycans

N-glycan analysis by lectin overlay was performed with lectins and carbo-free blocking solution obtained from Vector Laboratories (Kirtlington, UK) as per manufacturer’s instructions. N-glycan analysis by ultra-high performance liquid chromatography-mass spectrometry was essentially as described (44–46), see supplemental information for a full method.

### Immobilised immune complexes

96 wells were coated overnight at 4°C with 100μg/ml HSA in PBS, blocked with 1% fat-free milk in PBS for 1h, incubated with 2.5μg/ml (a)galactosylated human anti-HSA IgG1 in PBS for 2h at room temperature, and washed with PBS thrice before use.

### Neutrophil adhesion

Neutrophils were allowed to adhere to immobilised ICs in tissue culture grade 96 well plates as described (47). After fixation, non-overlapping brightfield images were taken of each well. Images were segmented using Cellpose3 (48), individual cells assigned an identity using the MorphoLibJ plug-in (49) in FiJi (National Institutes of Health, USA) and cell number, area, and eccentricity analyzed automatedly using an analysis pipeline in CellProfiler (The Broad Institute, Cambridge Massachusetts, USA).

### ROS production

ROS production was measured indirectly in a luminol assay using 5×10^5^ neutrophils per well in luminescence-grade 96 well plates coated with immobilised ICs in a Cytation plate reader (BioTek, Swindon, UK) as previously described (47).

### Comparison of FcγR expression neutrophils

FcγR phenotyping of healthy donor and patient neutrophil was performed using a spectral flow cytometer (Aurora, Cytek Biosciences, Fremont, California, USA). Spectral unmixing and data acquisition were performed using Spectroflo software (Cytek Biosciences).

### IgG binding

Agalactosylated and galactosylated IgG1 were incubated with PE-labelled F(ab’)_2_ fragments of goat anti-human Fab-specific fragments for 30 minutes at 37°C to form immune complexes (IgG-F’_2_) and added to 2×10^5^ CHO cells expressing individual neutrophil FcγRs as indicated exactly as previously described (20). IgG-F’_2_ IC binding was analyzed using flow cytometry (Attune NxT, Thermo Fisher Scientific). Flow cytometry data were analyzed using FCS Express (De Novo Software, Pasadena, USA).

### Statistical Analysis

Power calculations were not performed as part of this work. Statistical analysis was performed with GraphPad Prism 10 (GraphPad Software, Boston, USA). Where data met the assumptions for parametric tests, analysis was by two-tailed T test and by one-way ANOVA with Dunnet’s multiple comparison test were used for pairwise and multiple comparisons, respectively. Otherwise, nonparametric alternatives were used instead (Mann Whitney rank sum test and one-way ANOVA with Krustal-Wallis test, respectively). For kinetic ROS production experiments, the area under the curve was used for analysis. *p* values <0.05 were deemed statistically significant.

### Study Approval

Approval to use healthy donor blood was obtained from the Edinburgh Medical School Research Ethics Committee (21-EMREC-041 and 25-EMREC-023). Ethics to use patient samples was obtained via request SR2399 from the Lothian NHS Research Scotland Bioresource (25/ES/0030).

## Data availability

All data generated or analyzed during this study are included in this manuscript and its supplemental data files.

## Acknowledgements

We thank all patients and healthy volunteers who donated blood for this study. We thank the IRR flow facility staff for advice on flow cytometry and setting up spectral flow unmixing and the IRR imaging facility staff for advice on image analysis. We are grateful to Jenny Woof (University of Dundee) for helpful discussions and Graeme Cowan for advice on seamless cloning. pVITRO1-ΔV-IgG1/κ (Addgene plasmid #52213) was a gift from Andrew Beavil (King’s College, London).

## Author contributions

Experimentation – CP, AS, BF, JS; Data analysis - CP, AS, BF, JS, SV; Reagents and Advice – YN, HC, MC, PB, MG; Supervision – SV, CB, CH, GL; Funding acquisition – SV, HC; Wrote the draft – CP and SV; All authors edited and concur with the paper.

