## supplemental material for "Rheumatoid Arthritis-associated IgG N-glycan agalactosylation diminishes neutrophilic inflammation by reducing FcγR binding and downstream signaling"

**Supplemental Methods.**

*Analysis of glycans by ultra-high performance liquid chromatography-mass spectrometry****.***

IgG1 was denatured at 65°C with 1.33% sodium dodecyl sulfate (SDS) for 10 min. The inhibitory effect of SDS on PNGase F activity was neutralized with 4% Igepal, and N-glycans were enzymatically released with PNGase F (24 mU/µL in 5× PBS) at 37 °C for 18 h (44). Released glycans were fluorescently labeled with procainamide (43.2 mg/mL in dimethyl sulfoxide:acetic acid, 70:30, v/v) as described (46). This was via reductive amination in a two-step reaction using 2-picoline borane (44.8 mg/mL) as a reducing agent, with sequential incubations at 65 °C. Labeled glycans were purified by HILIC solid-phase extraction on a 0.2 µm AcroPrep wwPTFE filter plate, preconditioned with ethanol, water, and acetonitrile, and eluted with ultrapure water. N-glycans were concentrated using porous graphitized carbon (PGC) cleanup as described (45) with minor modifications. For this, samples were loaded onto a previously washed and equilibrated C18 ZipTip containing PGC in methanol suspension (50 mg/mL). The ZipTip was washed with acetonitrile and equilibrated with ultrapure water. Concentrated glycans were eluted with LC-MS grade acetonitrile:100 mM ammonium formate (pH 4.4; 75:25, v/v) by centrifugation at 2000 *g* for 30 s and analyzed by UHPLC coupled to ESI-qTOF-MS operating in positive ion mode. Procainamide-labeled N-glycans were separated on a Waters BEH Glycan column (100 × 2.1 mm, 1.7 µm) maintained at 60 °C with a flow rate of 0.4 mL/min. 100 mM ammonium formate (pH 4.4) was used as solvent A and acetonitrile as solvent B, applying a gradient of 75–62% B over 29 min within a 38 min analytical run, as described (44). Chromatograms were processed and manually integrated using Bruker DataAnalysis software. Results were expressed as percentage of the total normalized area. MS data were acquired over an m/z range of 50–1750 with a scan frequency of 0.5 Hz, and the three most intense precursor ions were selected for CID fragmentation. Features detected in MS spectra were searched for possible procainamide-labeled N-glycan compositions using the GlycoMod ExPASy tool (https://web.expasy.org/glycomod/) based on singly charged m/z values. Recorded MS/MS spectra were used to confirm annotated glycan structures, enabling structural assignment of individual chromatographic peaks.

**Supplemental Tables.**

| **Antibody** | **Clone** | **Isotype** | **Source** | **Identifier** |
| --- | --- | --- | --- | --- |
| APC-anti-CD16 | 3G8 | Mouse IgG1 | Biolegend | 302012 |
| PE-anti-CD32A | S20004A | Mouse IgG1 | Biolegend | 365504 |
| PE/Cy7-anti-CD32B/C | S18005H | Mouse IgG1 | Biolegend | 398314 |
| AF488-anti-CD64 | 10.1 | Mouse IgG1 | Biolegend | 305010 |
| APC/Cy7-anti-CD11b | M1/70 | Rat IgG2b | Biolegend | 101226 |
| Anti-phospho Akt | D25E6 | Rabbit IgG | CST | 13038 |
| Anti-Akt | 11E7 | Rabbit IgG | CST | 4685 |
| Anti-phospho Erk | D13.14.4E | Rabbit IgG | CST | 4370 |
| Anti-Erk | W15133B | Rat IgG1 | Biolegend | 686901 |
| Anti-phospho p38 | polyclonal | n/a | CST | 9211 |
| Anti-p38 | D13E1 | Rabbit IgG | CST | 8690 |
| HRP-anti-human IgG1 Fc | M1308A10 | Rat IgG2a | Biolegend | 410603 |
| Rabbit IgG | polyclonal | n/a | Sigma | I8140 |
| Rabbit Anti-HSA | polyclonal | n/a | Sigma | A0433 |
| HRP- anti-rabbit | polyclonal | n/a | BioRad | 170-6515 |

***Table S1. Details of commercially available antibodies used in this study.***

| **Inhibitor** | **Target** | **Concentration** | **Source** | **Identifier** |
| --- | --- | --- | --- | --- |
| BIRB-796 | pan-p38 | 100nM | Apex Biologix | A5639 |
| Ly2228820 | p38α, p38β | 500nM | Stemcell Technologies | 74162 |
| U-0216 | MEK1/2 | 10μM | Stemcell Technologies | 73522 |
| Ly294001 | pan-PI3K | 10μM | Synkinase | Syn-1108-M001 |
| DPI | NADPH oxidase | 10μM | Cayman Chemicals | 81050 |

***Table S2. Details of inhibitors used in this study.*** Concentration, final concentration used to inhibit targets. DPI, diphenyleneiodonium.

**Supplemental Figures.**

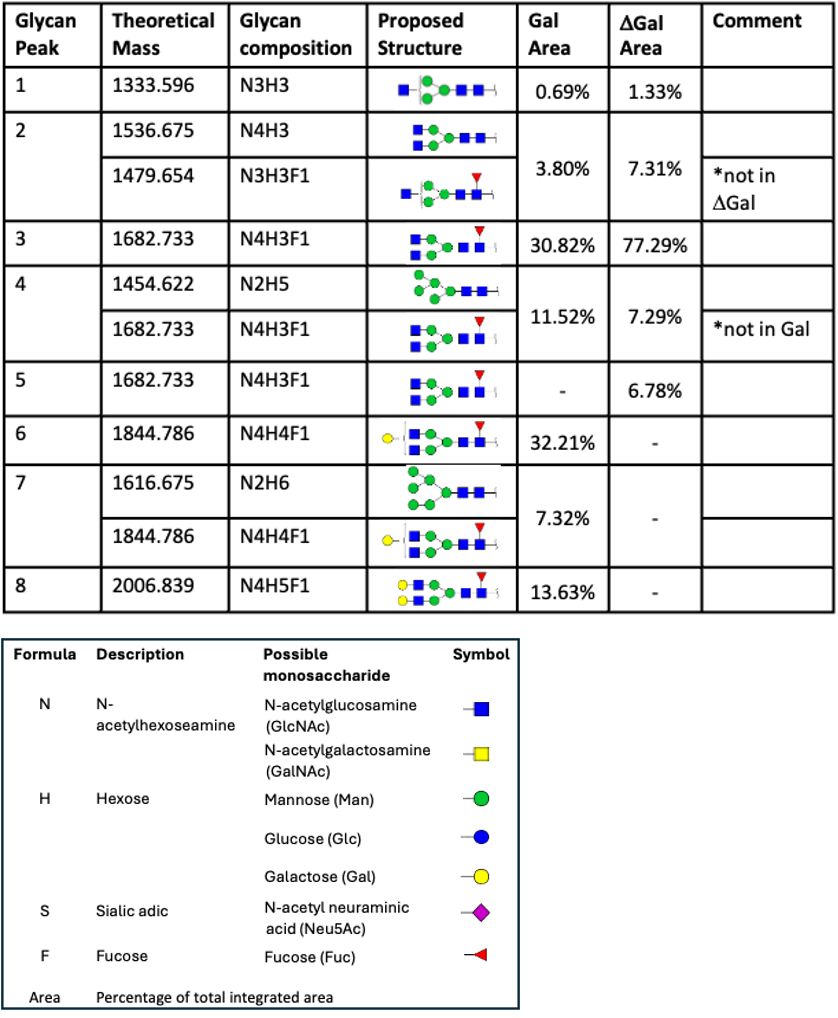
 ***Figure S1. Confirmation of glycans on recombinant IgG generated in glycoengineered CHO cells.*** Differences in relative abundance (% area) of procainamide-labelled IgG1 N-glycans between galactosylated (Gal) and agalactosylated (ΔGal) samples with corresponding N-glycan compositions and proposed structures for each peak (top) are presented alongside a legend (below).

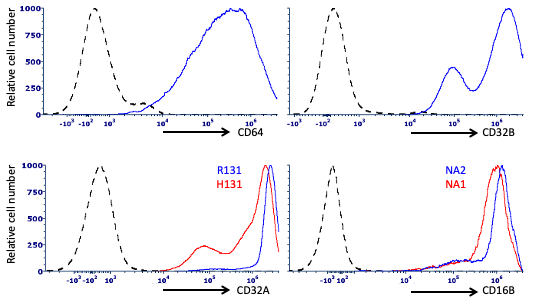
***Fig S2. Heterologous FcγR expression on CHO cells.*** FcγR expression on stably transfected CHO cells was analyzed by flow cytometry using antibodies listed in table S1. Black broken peaks represent untransfected cells. Continuous peaks represent CHO cells stably transfected with human (A) CD64/FcγRI; (B) CD32B/FcγRIIB; (C) CD32A/FcγRIIA (2 separate SNPs as indicated in blue and red); (D) CD16B/FcγRIIIB (2 separate SNPs as indicated in blue and red).
